# Generation of Recombinant Rotaviruses Expressing Human Norovirus Capsid Proteins

**DOI:** 10.1101/2022.08.11.503658

**Authors:** Asha A. Philip, John T. Patton

**Affiliations:** Department of Biology, Indiana University, Bloomington, IN 47405, USA

**Keywords:** rotavirus, reverse genetics, norovirus, vaccines, recombinant virus, plug-and-play expression vector, 2A-like translation element, viral expression vector

## Abstract

Rotaviruses, segmented dsRNA viruses of the *Reoviridae* family, are a primary cause of acute gastroenteritis in young children. In countries where rotavirus vaccines are widely used, norovirus (NoV) has emerged as the major cause of acute gastroenteritis. Towards the goal of creating a combined rotavirus-NoV vaccine, we explored the possibility of generating recombinant rotaviruses (rRVs) expressing all or portions of NoV GII.4 VP1 capsid proteins. This was accomplished by replacing the segment 7 NSP3 ORF with a cassette encoding sequentially NSP3, a 2A stop-restart translation element, and all or portions (P, P2) of NoV VP1. In addition to successfully recovering SA11 rRVs with modified SA11 segment 7 RNAs encoding NoV capsid proteins, analogous rRVs were recovered through modification of the RIX4414 segment 7 RNA. Immunoblot assay confirmed that rRVs expressed NoV capsid proteins as independent products. Moreover, VP1 expressed by rRVs underwent dimerization and was recognized by conformational-dependent anti-VP1 antibodies. Serially passaged rRVs that expressed the NoV P and P2 were genetically stable, retaining sequences up to 1.1 kbp without change. However, serially passaged rRVs containing the longer 1.5 kb VP1 sequence were less stable and gave rise to virus populations with segment 7 RNAs lacking VP1 coding sequences. Together, these studies suggest that it may be possible to develop combined rotavirus-NoV vaccines using modified segment 7 RNA to express NoV P or P2. In contrast, development of potential rotavirus-NoV vaccines expressing NoV VP1 will need additional efforts to improve genetic stability.

**Importance:** Rotavirus (RV) and norovirus (NoV) are the two most important causes of viral acute gastroenteritis (AGE) in young children. While the incidence of RV AGE has been brought under control in many countries through the introduction of live attenuated RV vaccines, similar highly effective NoV vaccines are not available. To pursue the development of a combined RV-NoV vaccine, we examined the potential usefulness of RV as an expression vector of all or portions of the NoV capsid protein VP1. Our results showed that by replacing the NSP3 open reading frame in RV genome segment 7 RNA with a coding cassette for NSP3, a 2A stop-restart translation element, and VP1, recombinant RVs can be generated that express NoV capsid proteins as separate products. These findings raise the possibility of developing a new generation of RV-based combination vaccines that can provide protection against a second enteric pathogen, such as the NoV.

## Introduction

Rotavirus (RV) and norovirus (NoV) are leading causes of severe acute gastroenteritis (AGE) in young children and the elderly (1,2). The introduction of effective RV vaccines - the RV1 monovalent vaccine, Rotarix (GSK Biologicals), and the RV5 pentavalent vaccine, RotaTeq (Merck and Company) – into the childhood immunization programs of the US and many other countries has resulted in significant reductions of the incidence of RV hospitalizations and mortality. The effectiveness of these vaccines has been correlated, in some instances, with the induction of neutralizing antibodies in immunized children (3-5). In countries where RV vaccines are widely used, NoV has emerged as the primary cause of diarrheal disease and diarrheal-associated hospitalizations in children during the first 5 years of life (6,7). The incidence of severe NoV disease is greatest in young children, in the elderly, and in subjects with compromised immunity (8). NoV is extremely contagious with fewer than 10 infectious particles able to cause AGE. The virus is highly stable in the environment and can be shed from individuals for weeks following infection (9). It has been difficult to develop effective anti-NoV therapeutics or vaccines due to the lack of suitable permissive cell lines and validated animal model systems (10,11).

Ten genogroups (GI-GX) and 49 genotypes of NoV have been identified based of sequence analysis of the NoV capsid protein VP1 (12). Human NoV disease is associated with strains belonging to genogroups I (GI), II (GII), IV (GIV), VII (GVII) or IX (GIX) (13). However, viruses of the GII genogroup are primarily responsible for human NoV disease worldwide; the GII.4 genotype has dominated since the 1990s and accounts for 55–85% of all NoV disease (14-16). A high degree of genetic diversity exists within each genogroup. Infections due to strains belonging to one genogroup generally do not confer protection against another genogroup, which challenges the development of universal NoV vaccines (17-18).

The NoV capsid is nonenveloped icosahedron comprised of 90 dimers of VP1 (Fig. 1) (11,18). The 7.6 kbp (+)RNA genome of NoV is organized into three open reading frames (ORF1-3). VP1 is the product of ORF3 and, in baculovirus expression systems, self-assembles into virus-like particles (VLPs). Based on its positioning in the NoV capsid, VP1 can be resolved into an interior shell (S) domain and a protruding (P) domain (19,20). Within genogroups, the S domain is relatively conserved in its primary sequence whereas the P domain is much more variable. The P domain can be further resolved into P1 and P2 subdomains. The highly variable P2 subdomain represents the immunodominant region of the VP1, serving as a target for neutralizing antibodies, and contains the receptor binding site for NoV capsid (21,22).

**Figure 1.**
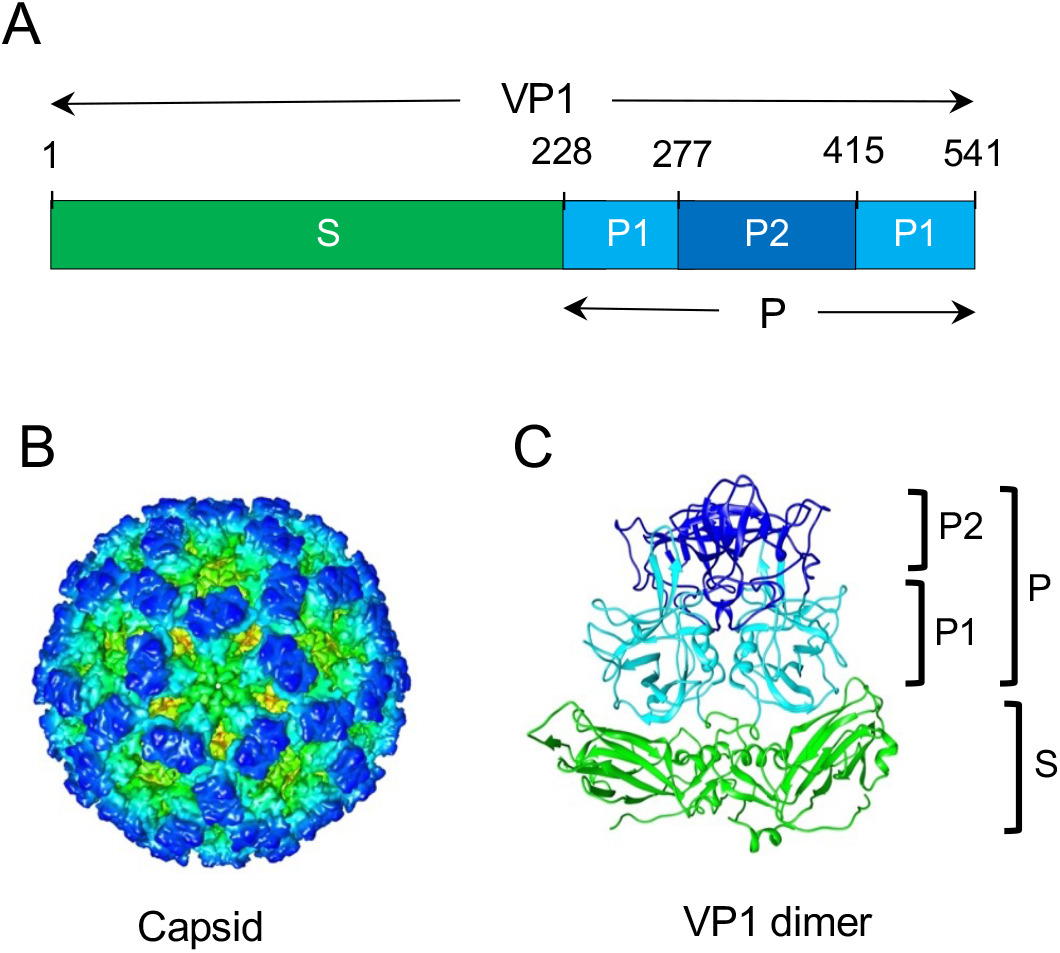
Domains of the human NoV VP1 capsid protein. **(A)** VP1 consists of shell (S) and protruding (P) domains. The P domain is further resolved into P1 and P2 subdomains. **(B)** Surface representation of NoV capsid with the S domain (green) and P1 (cyan) and P2 (blue) subdomains of VP1 distinguished by color. **(C**) Ribbon representation of a NoV VP1 dimer: S (green), P1 (cyan) and P2 (blue).

Three distinct types of NoV vaccine candidate have been developed to date: NoV VLPs and P particles, and recombinant adenoviruses expressing NoV capsid proteins (23,24). In general, studies evaluating NoV vaccine candidates have been performed in adults (25-27). However, a phase II trial of a potential GI.1 and GII.4 bivalent VLP vaccine carried out in children and infants showed that the candidate was safe and evoked a robust immune response (28). In an attempt to simultaneously prevent both RV and NoV diseases, a trivalent vaccine containing two types of NoV GII.4 VLPs (GII.4 NO-2010 and GII.4 SYD-2012) and oligomeric RV VP6 was tested in a mouse model (29,30). The results demonstrated significant enhancement in the production of cross-reactive anti-VP1 IgG antibodies following co-immunization of GII.4 VLPs and VP6 (28). RV VP6 is a highly conserved protein, capable of evoking a protective anti-RV immune response in immunized animals, and can act as an adjuvant stimulating immune responses to NoV antigen (31,32).

Recent development of RV reverse genetics systems has resulted in the generation of recombinant RVs that can serve as expression vectors of foreign proteins (33-40). The RV genome consists of 11 segments of dsRNA, with a total size of 18.5 kbp (1). All the segments contain a single ORF except for segment 11, which contains two out-of-frame ORFs. Together, the genome segments encode six structural (VP) and six non-structural (NSP) viral proteins (1). Genome segment 7 of group A rotavirus encodes the 36 kDa protein, NSP3, an RNA-binding protein that acts as a translation enhancer of viral plus-sense (+) RNAs in infected cells (41-43). In our previous studies, we showed that segment 7 can be engineered to express NSP3 and a separate foreign protein by inserting a 2A stop-restart translational element immediately downstream of the NSP3 ORF followed by the ORF for the foreign protein (36,37). This approach resulted in the production of recombinant RVs expressing fluorescent proteins (FPs) [UnaG (green), mRuby (red), Tag BFP (blue)] from segment 7 (37). Similarly, as a step in exploring the potential of RV as a vaccine expression vector, we showed that segment 7 could be modified to express, as separate products, portions of the SARS-CoV-2 spike protein using 2A translation elements (38). Analysis of RV expressing foreign proteins showed that they are generally well growing and genetically stable and continue to produce functional NSP3, capable of dimerization and inducing nuclear localization of the cellular poly(A)-binding protein (PABP) (36,37). These results raise the possibility of engineering RVs into vector systems that express NoV capsid proteins, enabling the production of combined RV-NoV vaccines that may provide immunological protection against the two most common causes of childhood AGE. In this study, we show that it is possible to generate recombinant RVs that express all or portions (P, P2) of the VP1 capsid protein of a human GII.4 NoV.

## Results

### Construction of modified segment 7 plasmids encoding NoV capsid proteins

To evaluate the possibility of using recombinant rotaviruses as expression vectors of NoV capsid proteins, we replaced the NSP3 ORF in pT7/NSP3SA11 with a cassette that sequentially encoded NSP3, a flexible linker (GAG), a porcine teschovirus 2A translational stop-restart element, and the NoV P2, P or VP1 protein (Fig. 2). Some cassettes included sequences specifying a 3xFLAG tag at the N terminus or 1xFLAG tag at the C terminus of the NoV capsid protein (Fig. 2). Instead of FLAG tags, some cassettes encoded NoV capsid proteins with a C-terminal 6xHis tag, with or without an intervening thrombin (Th) cleavage site (Fig. 2). This process yielded a set of pT7/SA11NSP3-2A-NoV vectors designed to express separately NSP3 and FLAG- or His-tagged P2 (pT7/NSP3-2A-fP2), P (pT7/NSP3-2A-fP or pT7/NSP3-2A-PHis), or VP1 (pT7/NSP3-2A-fVP1, pT7/NSP3-2A-VP1f, pT7/NSP3-2A-VP1-ThHis, or pT7/NSP3-2A-VP1 His). The NoV capsid sequences were introduced into the pT7/NSP3SA11 vector at the same site used previously for making rSA11 viruses that expressed epifluorescent reporter proteins and domains of the SARS CoV-2 spike protein (37,38).

**Figure 2.**
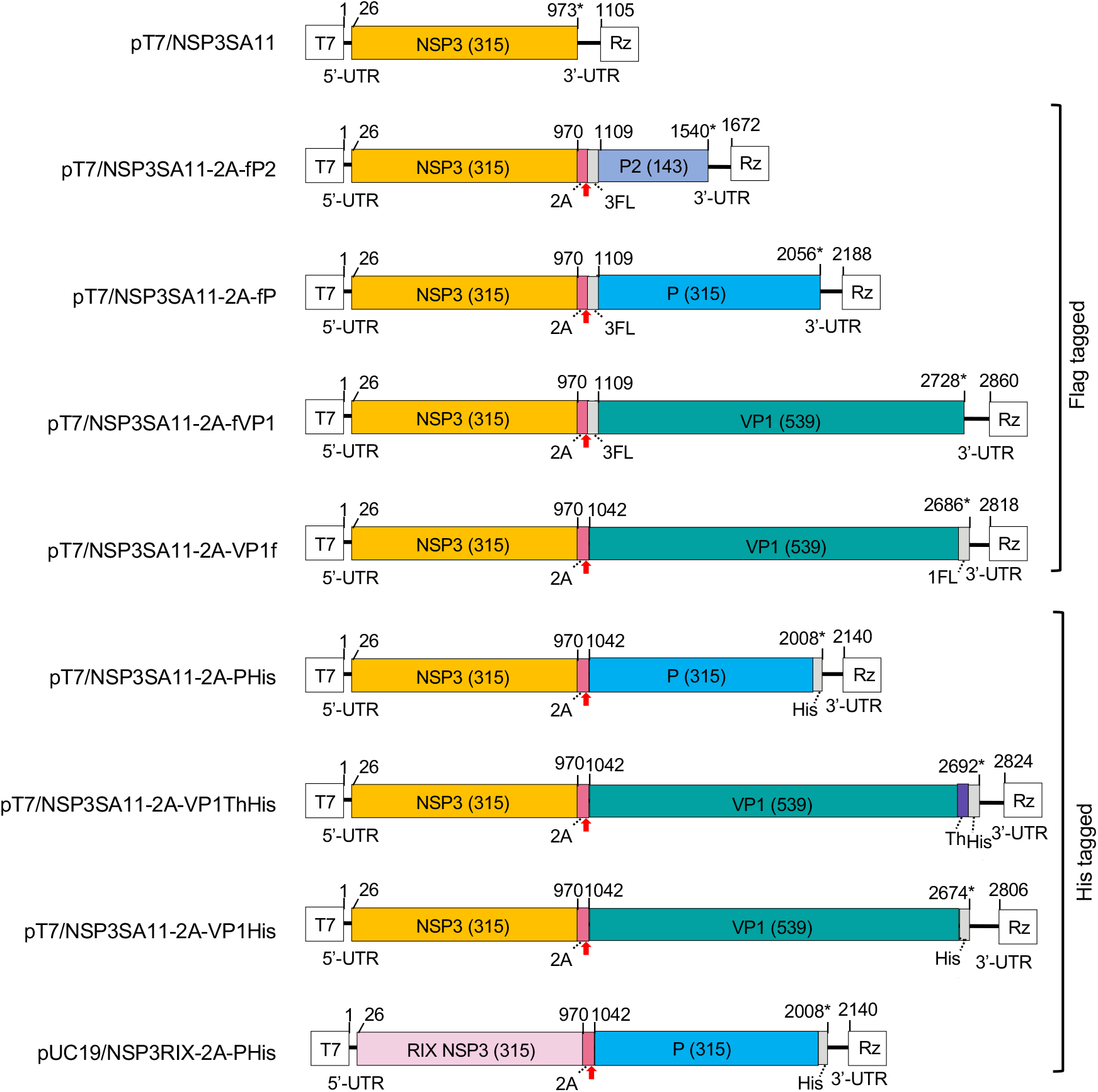
Plasmids with modified segment 7 cDNAs used to generate recombinant rotaviruses expressing NoV capsid proteins. Illustration indicates nucleotide positions of the coding sequences for NSP3, 2A element (2A), 3xFLAG (3FL), 1xFLAG (1FL), 6x histidine (His) tag, thrombin cleavage site (Th), and VP1, P2, and P. The red arrow notes the position of the 2A translational stop-restart site, and the asterisk notes the end of the ORF. Sizes (aa) of encoded NSP3 and NoV VP1 products are given in parenthesis. T7 (T7 RNA polymerase promoter sequence), Rz (hepatitis D virus ribozyme), UTR (untranslated region).

To test the possibility that segment 7 RNAs of RV strains, besides SA11, could serve as expression platforms, we generated a RIX4414-like (RIX)-based segment 7 plasmid (pUC19/NSP3RIX-2A-PHis) in which the NSP3 ORF had been replaced with a coding cassette for NSP3-GAG-2A-NoV P-6xHis (Fig. 2). RIX4414 is a human RV G1P[8] strain that is used in formulating the Rotarix vaccine.

### Recovery of recombinant rotaviruses containing NoV coding sequences

Recombinant viruses were generated by transfecting BHK-T7 cells with a complete set of RV pT7 plasmids, each directing the synthesis of one of the eleven viral +RNAs, and a CMV expression vector for the African swine fever virus capping enzyme (NP8688R). In the transfection procedure, the pT7/NSP3SA11 plasmid was replaced with a pT7/SA11NSP3-2A-NoV or pT7/NSP3RIX-2A-PHis plasmid. The pT7/NSP2SA11 and pT7/NSP5SA11 plasmids were included in transfection mixtures at levels 3-fold greater than the other plasmids. Transfected BHK-T7 cells were over-seeded with MA104 cells 2 days post transfection. Three days later, cell mixtures were freeze-thawed, and recombinant viruses in the lysates were amplified by passage on MA104 cells. Recombinant viruses were recovered from virus pools by plaque selection and amplified to larger volumes prior to characterization. Properties of recombinant viruses are summarized in Table 1.

**Table 1.**
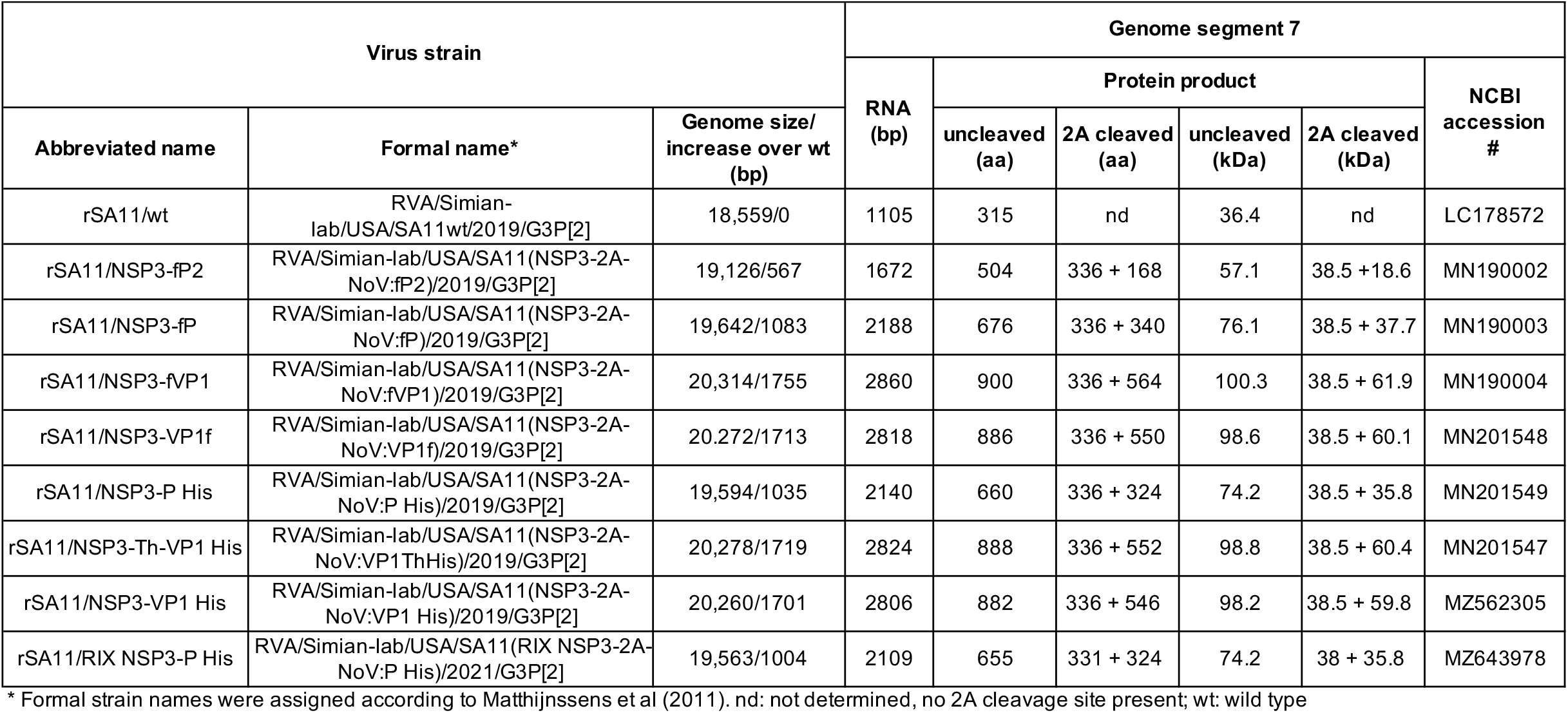
Properties of recombinant rRV/NSP3-2A-NoV strains.

| Virus strain |  |  | Genome segment 7 |  |  |  |  |  |
| --- | --- | --- | --- | --- | --- | --- | --- | --- |
|  |  |  | RNA (bp) | Protein product |  |  |  | NCBI accession # |
| Abbreviated name | Formal name* | Genome size/ increase over wt (bp) |  | uncleaved (aa) | 2A cleaved (aa) | uncleaved (kDa) | 2A cleaved (kDa) |  |
| rSA11/wt | RVA/Simian-lab/USA/SA11wt/2019/G3P[2] | 18,559/0 | 1105 | 315 | nd | 36.4 | nd | LC178572 |
| rSA11/NSP3-fP2 | RVA/Simian-lab/USA/SA11(NSP3-2A-NoV:fP2)/2019/G3P[2] | 19,126/567 | 1672 | 504 | 336 + 168 | 57.1 | 38.5 + 18.6 | MN190002 |
| rSA11/NSP3-fP | RVA/Simian-lab/USA/SA11(NSP3-2A-NoV:fP)/2019/G3P[2] | 19,642/1083 | 2188 | 676 | 336 + 340 | 76.1 | 38.5 + 37.7 | MN190003 |
| rSA11/NSP3-fVP1 | RVA/Simian-lab/USA/SA11(NSP3-2A-NoV:fVP1)/2019/G3P[2] | 20,314/1755 | 2860 | 900 | 336 + 564 | 100.3 | 38.5 + 61.9 | MN190004 |
| rSA11/NSP3-VP1f | RVA/Simian-lab/USA/SA11(NSP3-2A-NoV:VP1f)/2019/G3P[2] | 20,272/1713 | 2818 | 886 | 336 + 550 | 98.6 | 38.5 + 60.1 | MN201548 |
| rSA11/NSP3-P His | RVA/Simian-lab/USA/SA11(NSP3-2A-NoV:P His)/2019/G3P[2] | 19,594/1035 | 2140 | 660 | 336 + 324 | 74.2 | 38.5 + 35.8 | MN201549 |
| rSA11/NSP3-Th-VP1 His | RVA/Simian-lab/USA/SA11(NSP3-2A-NoV:VP1ThHis)/2019/G3P[2] | 20,278/1719 | 2824 | 888 | 336 + 552 | 98.8 | 38.5 + 60.4 | MN201547 |
| rSA11/NSP3-VP1 His | RVA/Simian-lab/USA/SA11(NSP3-2A-NoV:VP1 His)/2019/G3P[2] | 20,260/1701 | 2806 | 882 | 336 + 546 | 98.2 | 38.5 + 59.8 | MZ562305 |
| rSA11/RIX NSP3-P His | RVA/Simian-lab/USA/SA11(RIX NSP3-2A-NoV:P His)/2021/G3P[2] | 19,563/1004 | 2109 | 655 | 331 + 324 | 74.2 | 38 + 35.8 | MZ643978 |

\* Formal strain names were assigned according to Matthijnssens et al (2011). nd: not determined, no 2A cleavage site present; wt: wild type

As expected, rSA11 viruses generated with pT7/NSP3-2A-NoV plasmids (subsequently referred to as rSA11/NSP3-2A-NoV viruses) contained segment 7 RNAs that were much larger than the segment 7 RNA of wild type SA11 virus (Fig. 3A and 4A). Sequence analysis confirmed that the segment 7 RNAs of the rSA11/NSP3-2A-NoV viruses matched those of the pT7/NSP3-2A-NoV plasmids. The addition of NoV P2 and P sequences into the 1.1-kbp segment 7 RNAs increased their total size to 1.7 kbp and 2.1 kbp, respectively (Table 1, Fig. 3A and 4A). Likewise, addition of NoV VP1 sequences to the segment 7 RNA increased its total size to 2.8-2.9 kbp, which migrated electrophoretically near genome segment 1. Insertion of NoV VP1 sequences into the segment 7 RNA increased the total size of the viral genome to 20.3 kbp, ∼9.5% greater than the size of the wildtype SA11 genome. The 2.8-2.9 kbp segment 7 RNAs of rSA11 viruses containing the NoV VP1 sequences are smaller than the 3.3 kbp segment 7 RNA of the rSA11/NSP3-fS1 virus, which contains the SARS CoV-2 S1 coding sequence (38).

**Figure 3.**
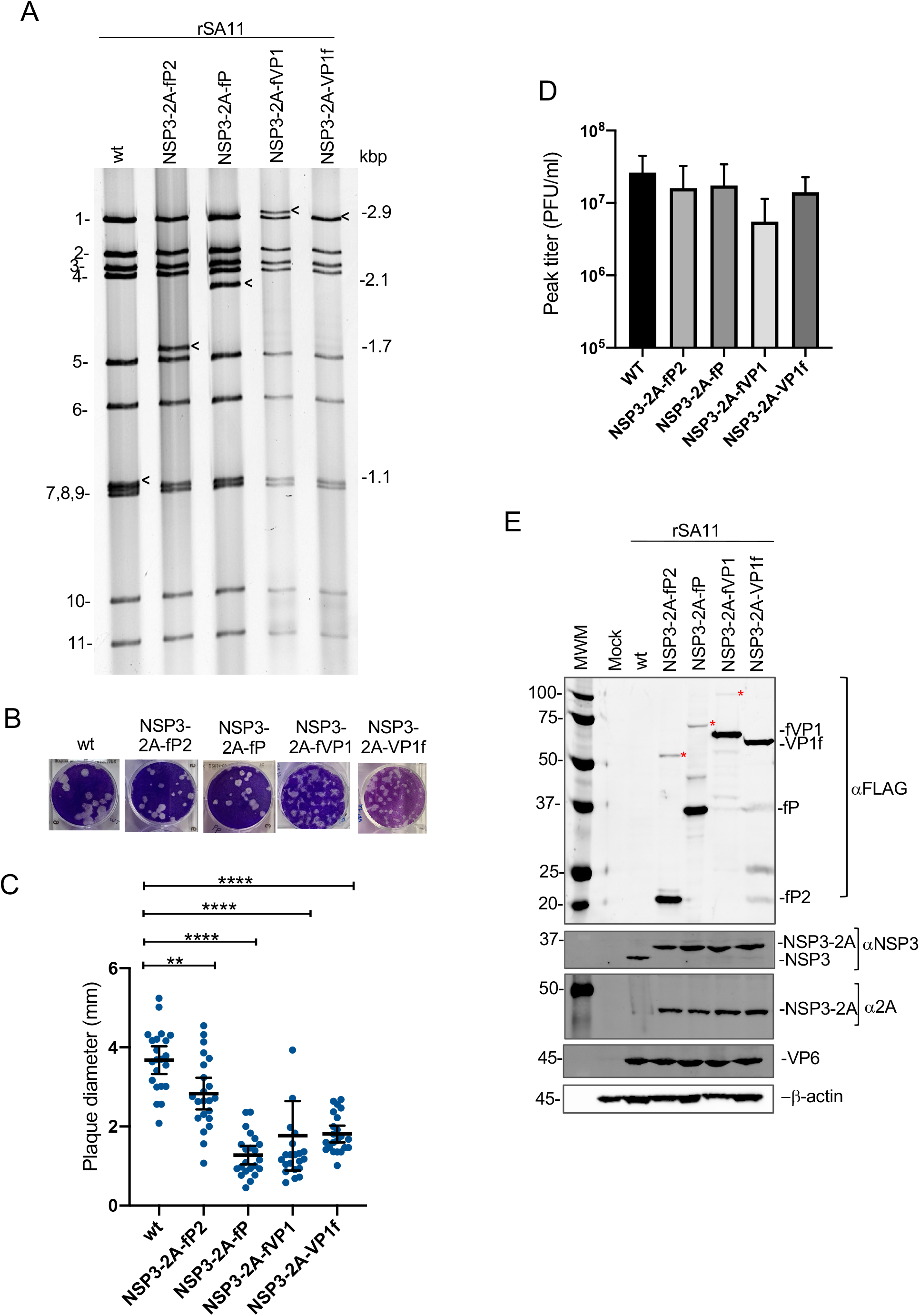
rSA11 viruses expressing FLAG-tagged NoV capsid proteins. **(A)** dsRNAs were recovered from rSA11-infected MA104 cells, resolved by gel electrophoresis, and detected by ethidium-bromide staining. The genome segments of rSA11/wt are labeled 1 to 11. Sizes (kbp) of modified segment 7 RNAs (black arrows) are indicated. **(B)** Plaques were detected by crystal-violet staining. **(C)** Mean diameters of rSA11 plaques at 5 days p.i., with 95% confidence intervals indicated (black lines). **(D)** Titers reached by rSA11 isolates were determined by plaque assay (plaque-forming units, PFU). **(E)** Immunoblot assay was used to examine mock and rSA11-infected cells for the presence of NoV VP1, P, or P2 (αFLAG antibody), NSP3 or NSP3-2A (αNSP3), NSP3-2A (α2A), VP6, and β-actin. Read through products, resulting from lack of 2A peptide function, are indicated with a red asterisk. Sizes (kDa) of protein molecular weight markers (MWM) are indicated.

**Figure 4.**
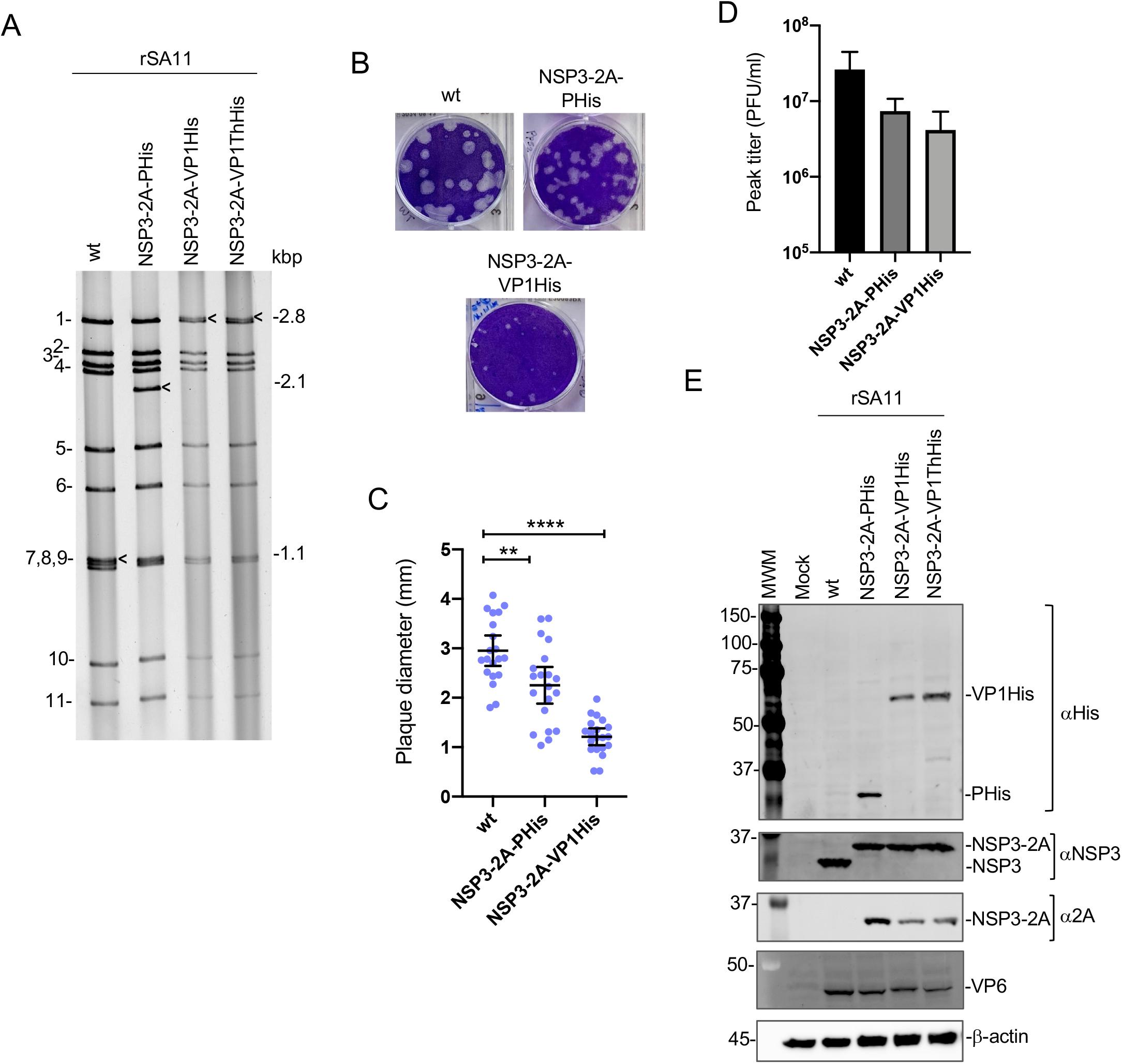
rSA11 viruses expressing His-tagged NoV capsid proteins. **(A)** dsRNAs were recovered from rSA11-infected MA104 cells, resolved by gel electrophoresis, and detected by ethidium-bromide staining. RNA segments of rSA11/wt are labeled 1 to 11. Sizes (kbp) of modified segment 7 RNAs (black arrows) are indicated. **(B)** Viral plaques assays were detected by crystal-violet staining. **(C)** Mean diameters of plaques at 5 days p.i., with 95% confidence intervals indicated (black lines). **(D)** Titers reached by rSA11 isolates were determined by plaque assay (plaque-forming units, PFU). **(E)** Immunoblot assay was used to examine mock and rSA11-infected cells for the presence of NoV PHis, VP1His, and VP1ThHis (α6xHis antibody), NSP3 or NSP3-2A (αNSP3), NSP3-2A (α2A), VP6, and β-actin. Sizes (kDa) of protein molecular weight markers (MWM) are indicated.

Analysis of recombinant viruses recovered using the pT7/NSP3RIX-2A-PHis plasmid confirmed that it was possible to use the RIX segment 7 RNA to produce viruses containing coding sequences for the NoV capsid protein (Fig. 5A). The rSA11/(RIX)NSP3-2A-NoV PHis viruses contain a 2.1 kbp segment 7 RNA instead of the wildtype 1.1 kbp segment 7 RNA.

Plaque analysis showed that the plaques formed by rSA11/NSP3-2A-NoV and rSA11/(RIX)NSP3-2A-NoV P viruses were smaller than those of rSA11/wt virus (Fig. 3B,C, 4B,C and 5B). This is consistent with previous studies showing that rRVs with modified segment 7 RNAs expressing FPs or SARS-CoV-2 proteins have a small plaque phenotype (37,38). Plaque analysis also indicated that rSA11/NSP3-2A-NoV viruses grew to maximum titers in MA104 cells that were ≥1/2 log less than rSA11/wt (Fig. 3D, 4D, and 5C). The basis for the smaller plaque phenotypes and lower titers is unknown but could be due to a longer elongation time required for the viral RNA polymerase to transcribe the modified segment 7 dsRNAs or a longer time required for translating the segment 7 mRNA containing foreign protein sequences. Alternatively, it may reflect the complexity associated with packaging the large modified dsRNA containing foreign sequences and assembly of the viral particles.

### Recombinant viruses express NoV P2, P or VP1 as separate products

To examine NoV protein products made by rSA11/NSP3-2A-NoV and rSA11/(RIX)NSP3-2A-NoV P viruses, lysates prepared at 9 h p.i. from MA104 infected cells were probed by immunoblot assay using FLAG antibody (Fig. 3E). The anti-FLAG immunoblots showed that rSA11/NSP3-2A-fP2, -fP, -fVP1 and -VP1f viruses directed the expression of NoV proteins of the size predicted for a functional 2A element in the modified NSP3 ORF: -fP2 (18.6 kDa), -fP (37.7 kDa), -fVP1 (61.9 kDa) and -VP1f (60.1 kDa) (Table 1). Assays with anti-NSP3 antibody and 2A element antibody identified a 38 kD protein, which corresponds to NSP3 (36 kD) linked to remnant residues of the 2A peptide (2 kDa) (37,44). Anti-FLAG immunoblots assay also detected minor amounts of large proteins, which likely represent read-through products of NSP3-2A-NoV protein cassettes (NSP3-2A-fP2, NSP3-2A-fP and NSP3-2A-fVP1) generated when the 2A element fails to function (Fig. 3E, red asterisks). Such read-through product was not detected for the rSA11/NSP3-2A-VP1f virus, suggesting that direct fusion of the VP1 ORF to the upstream 2A peptide sequence perhaps increases the efficiency of 2A activity (Fig. 3E).

Immunoblot assays performed with antibody specific for a 6xHis epitope showed that the rSA11/NSP3-2A-PHis virus, and the rSA11/NSP3-2A-VP1His and rSA11/NSP3-2A-VP1ThHis viruses generated 36 kDa P and 60 kDa VP1 products, respectively, sizes consistent with the presence of functional 2A peptides in the expression cassettes (Table 1, Fig. 4E). As noted above for rSA11/NSP3-2A-VP1f, the 6xHis antibody did not detect readthrough products (NSP3-2A-PHis, NSP3-2A-VP1His, and NSP3-2A-PThHis), possibly because placement of NoV P and VP1 coding sequences immediately downstream of the 2A peptide sequence, without intervening tags or spacer sequences, may improve the efficiency of 2A activity. Mirroring results described above for viruses expressing FLAG-tagged NoV protein products, immunoblot assays performed with antibodies for NSP3 and the 2A peptide revealed the expression of the 38kDa NSP3-2A protein in MA104 cells infected with rSA11/NSP3-2A-PHis, -VP1His, and -VP1ThHis (Fig. 4E).

Immunoblot assays with the anti-6xHis antibody showed that the 36 kDa P protein was a dominant product of rSA11/(RIX)NSP3-2A-PHis virus (Fig. 5D). Lesser amounts of a NSP3-2A-PHis read-through product was also detected with the anti-6xHis antibody. Due to lack of cross-reactivity, immunoblots probes with antibody raised against SA11 NSP3 failed to detect RIX NSP3 (Fig. 5D). However, re-probing the same blot with antibody specific for the 2A peptide identified expression of 38 kDa protein, likely representing RIX NSP3 protein fused to 2A peptide (Fig. 5D). Overall, these results show that it is possible to use segment 7 RNAs of rotavirus strains other than SA11, including the vaccine RIX4414 strain, to direct the expression of NoV capsid proteins.

**Figure 5.**
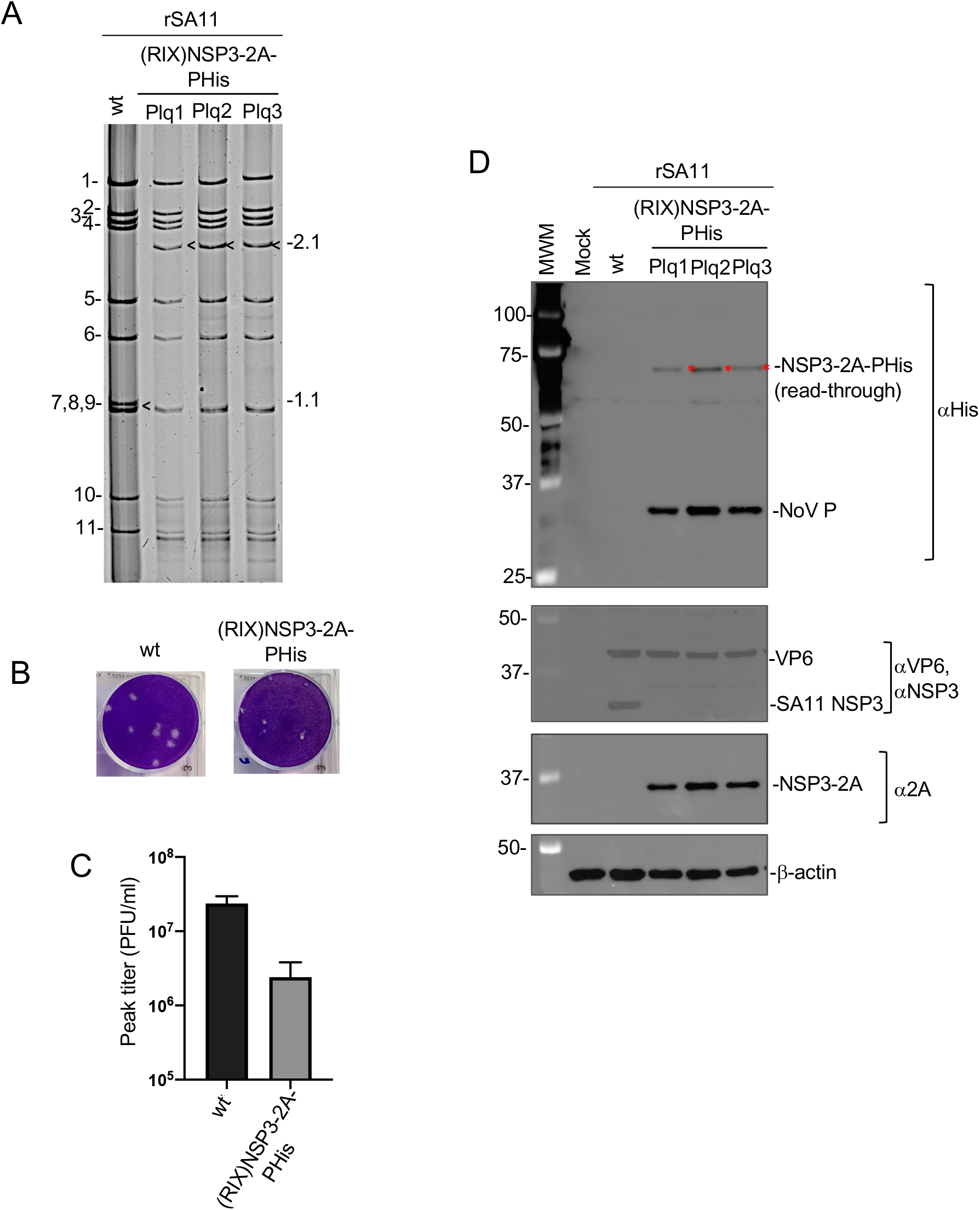
Recombinant virus expressing NoV P from modified RIX segment 7 RNA. **(A)** dsRNA was recovered from MA104 cells infected with rSA11 or a plaque isolate (Plq1-Plq3) of rSA11/NSP3RIX-2A-PHis, resolved by gel electrophoresis, and detected by ethidium-bromide staining. RNA segments of rSA11/wt are labeled 1 to 11. Size (kbp) of segment 7 RNAs (black arrows) are indicated. **(B)** Viral plaques assays were detected at 5 days p.i. by crystal-violet staining. **(C)** Titers reached by rSA11 isolates were determined by plaque assay. **(D)** Immunoblot assay was used to examine mock and rSA11-infected cells for the presence of PHis (α6xHis antibody), NSP3 or NSP3-2A (αNSP3), NSP3-2A (α2A), VP6, and β-actin. Read through products, resulting from lack of 2A peptide function, are indicated with a red asterisk. Sizes (kDa) of protein molecular weight markers (MWM) are indicated.

### Dimerization of the NoV VP1 expressed by recombinant viruses

VP1 dimers serve as intermediates in the assembly of the NoV capsid (45). To address whether NoV proteins expressed from recombinant viruses formed dimers, lysates from rSA11/NSP3-2A-fP2, -fP, - fVP1, and -VP1f infected cells were incubated sample buffer at temperatures in which VP1 dimers are stable (25°C) and temperatures in which they are disrupted (95°C). The samples were analyzed by gel electrophoresis and immunoblot assay using anti-FLAG antibody. The results indicated that neither expressed P2 nor P formed stable dimers. In contrast, expressed VP1 proteins, with FLAG tags at the N- or C-terminus (fVP1 and VP1f), migrated with the expected size of VP1 dimers (Fig. 6A). Under the same electrophoretic conditions used to analyze NoV capsid proteins for dimerization, cells infected with rSA11/wt and rSA11/NSP3-2A-fP2, -fP, -fVP1 and -VP1f generated NSP3 dimers and VP6 trimers (Fig. 6A), naturally occurring multimeric forms of these proteins (46,47). Similarly, 6xHis-tagged VP1 proteins (VP1His and VP1ThHis) expressed by rSA11/NSP3-2A-VP1His and rSA11/NSP3-2A-VP1ThHis also formed dimers (Fig. 6B). As above, NSP3 and VP6 proteins formed stable dimers and trimers, respectively, in rSA11/NSP3-2A-VP1His and rSA11/NSP3-2A-VP1ThHis infected cells. (Fig. 6B).

**Figure 6.**
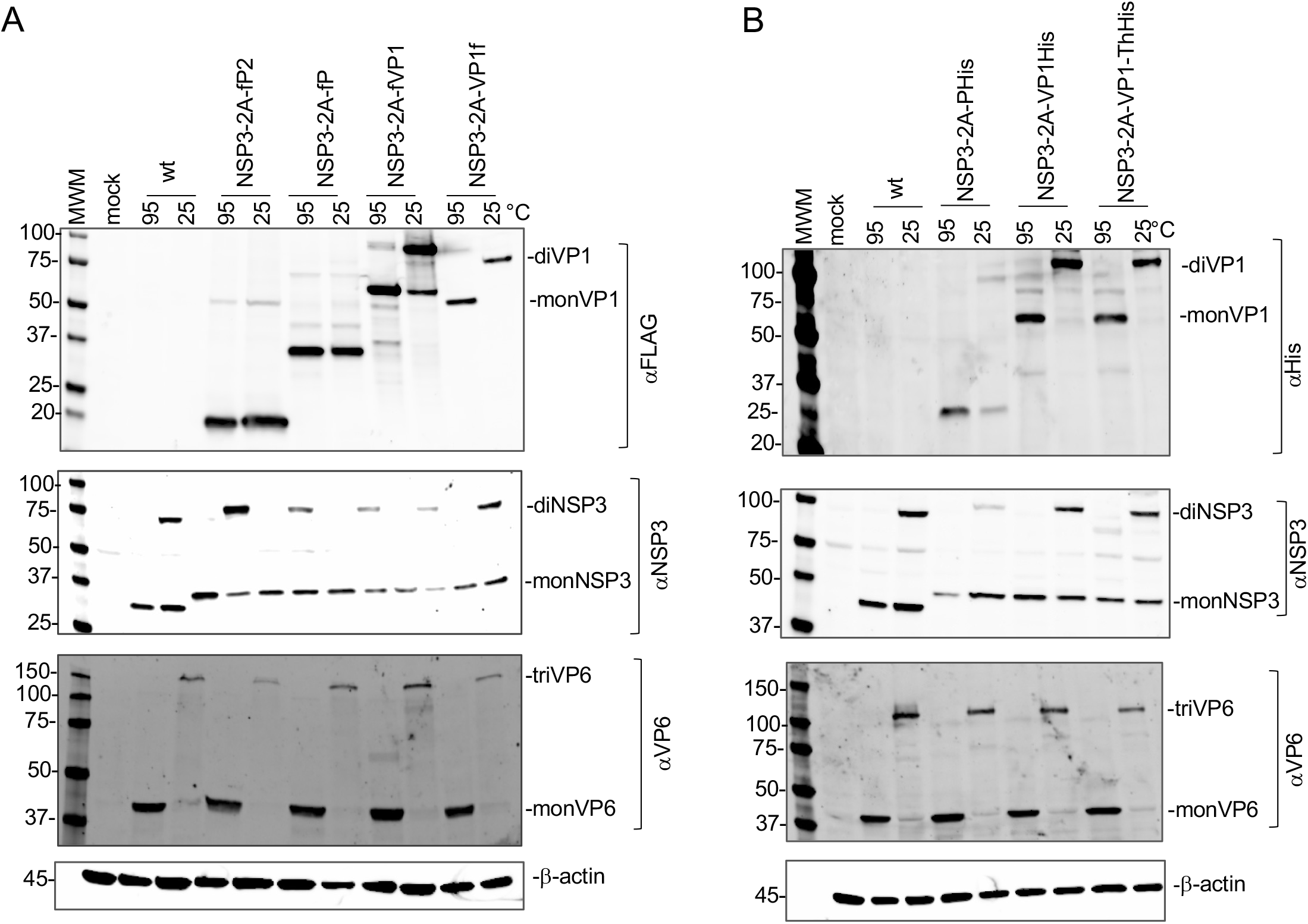
Dimerization of NoV capsid proteins expressed by rSA11. **(A)** MA104 cells were mock infected or infected with rSA11/wt or rSA11/NSP3-2A-fP2, -fP, -fVP1 or -VP1f and harvested at 9 h p.i. Cell lysates were mixed with SDS-sample buffer, incubated for 10 min at 25 or 95!C, and subjected to electrophoresis on a 4 to 20% polyacrylamide gels. Resolved proteins were blotted onto nitrocellulose membranes and detected using antibodies specific for FLAG, SA11 NSP3, VP6, or β-actin. Primary antibodies were detected using HRP-conjugated secondary antibodies. Sizes (kilodaltons) of protein markers (MWM) are indicated. **(B)** As in **(A)**, except cells were infected with rSA11/wt or rSA11/NSP3-2A-PHis, VP1His, or -VP1ThHis. Blots were probed with antibodies specific for 6xHis instead of FLAG.

### Folding of NoV VP1 capsid proteins into native structures

To gain insight into whether the VP1 products expressed from rSA11/NSP3-2A-VP1 viruses folded into native structures, lysates prepared from MA104 cells infected with rSA11/NSP3-2A-fVP1, -VP1f, and - VP1His viruses were probed by pulldown assay using a conformation-dependent neutralizing monoclonal antibody that recognizes GII.4 VP1 (NVB43.9). As shown in Fig. 7 (red arrows), the anti-VP1 antibody immunoprecipitated both FLAG and 6xHis tagged NoV VP1 proteins (fVP1 and VP1His), indicating that VP1 expressed by rSA11 viruses folded in a conformation that included an authentic neutralizing epitope found in the NoV VP1 protein. Combined with data indicating that rSA11-expressed VP1 formed dimers (Fig. 6), these data suggest at least some VP1 produced in cells infected with rSA11/NSP3-2A-fVP1 and -VP1His folded into native structures. Unlike the successful pulldown of fVP1 and VP1His with anti-GII.4 VP1 antibody, it was not certain whether the antibody likewise immunoprecipitated the VP1f product of rSA11/NSP3-2A-VP1f (Fig. 7). This may in part result from lower level of VP1f expression in infected cells as compared to fVP1 and VP1His or to differences in the affinity the FLAG antibody for the N-terminal 3xFLAG tag for fVP1 versus the C-terminal 1xFLAG tag for VP1f (Fig. 7).

**Figure 7.**
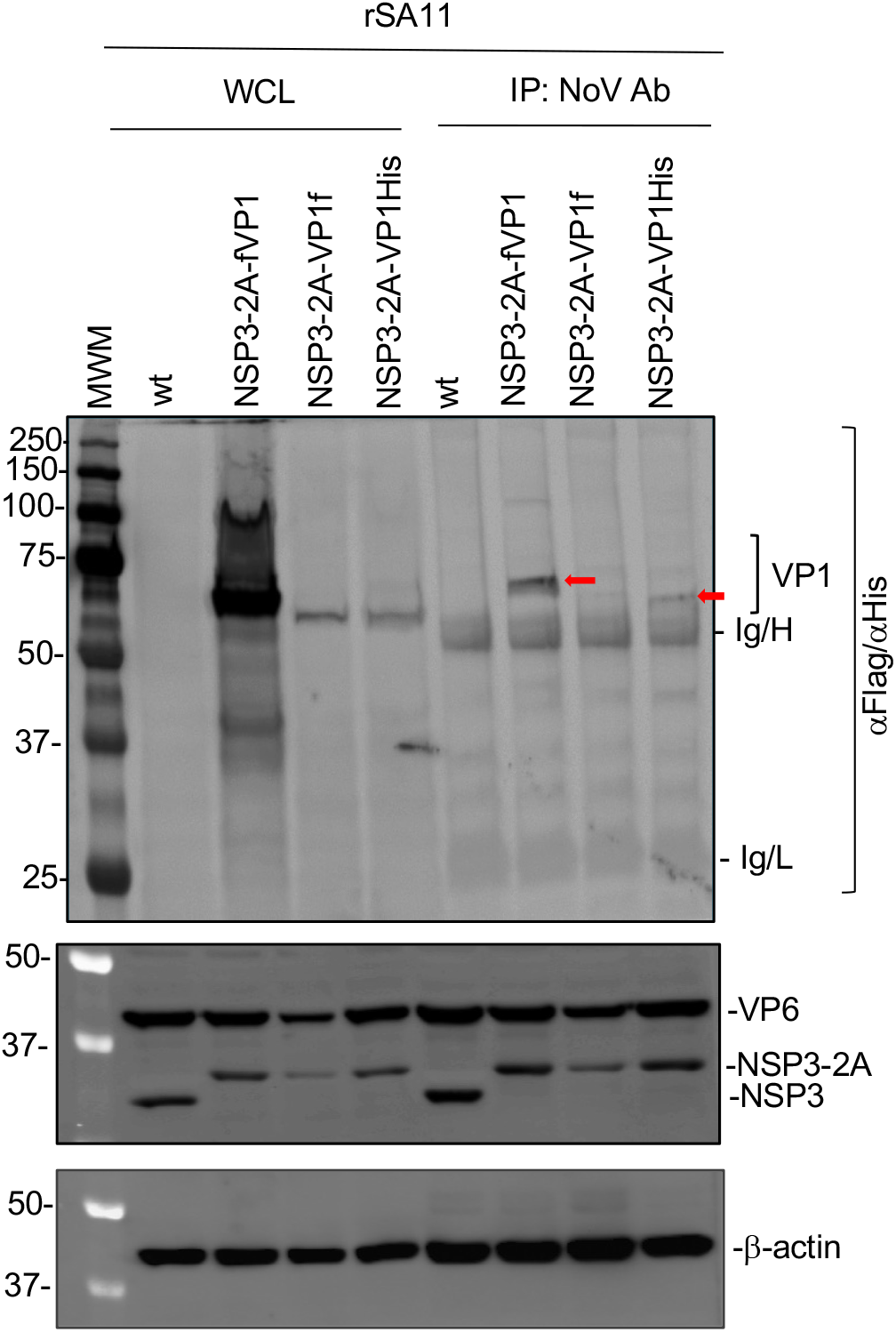
Recognition of a neutralizing conformationally-dependent epitope in NoV VP1 expressed by rSA11. Whole cell lysates (WCL) were prepared from MA104 cells infected with the indicated recombinant viruses (rSA11/wt; rSA11/NSP3-2A-fVP1, -VP1f, and -VP1His) and incubated with a monoclonal antibody specific for NoV GII.4 VP1 (NVB43.9). Antigen-antibody complexes were recovered using magnetic IgA/G beads, resolved along with whole cells lysates by gel electrophoresis, and blotted onto nitrocellulose membranes. Blots were probed with anti-FLAG/6xHis antibodies to detect immunoprecipitated VP1 and with anti-NSP3, -VP6, and –β-actin antibodies. Arrows indicate immunoprecipitated VP1 proteins. Ig light chain, (Ig/L) and Ig heavy chain, Ig/H. Positions of molecular weight markers (MWM) in kDa are indicated.

### Genetic stability of rSA11 strains expressing NoV proteins

To analyze the genetic stability of rSA11 viruses expressing NoV capsid proteins, the viruses were subjected to 5 rounds of serial passage at three dilutions (1:10, 1:100 or 1:1000). Analysis of the dsRNAs recovered from cells infected with the rSA11/NSP3-2A-fP2, -fP, fHis viruses showed no changes in the sizes of any of the genome segments, including the modified genome segment 7 RNA, over 5 rounds of serial passage (P1 to P5) (Fig. 8A). Thus, recombinant viruses carrying up to 1.1 kbp of foreign sequence were genetically stable, consistent with the findings of a previous study (37,38). In contrast, serial passage of rSA11/NSP3-2A-fVP1 and -VP1His showed evidence of genetic instability during serial passage at all three dilutions (Fig. 8B, C). Notably, new genome segments were visible by the third round of passage that were smaller (Fig. 8B, C; red arrows) than the initial 2.8-2.9 kbp segment 7 RNAs of NSP3-2A-fVP1 and -VP1His (black arrow heads). Indeed, the 2.8-2.9 kbp segment 7 RNA of these viruses were no longer detectable by passage 5. Instead, the ∼1.2 kbp variant RNA became a dominant genome segment for rSA11/NSP3-2A-fVP1 and -VP1His by passage 5, replacing the 2.8-2.9 kbp segment 7 RNA (Fig. 8B, C; red arrows).

**Figure 8.**
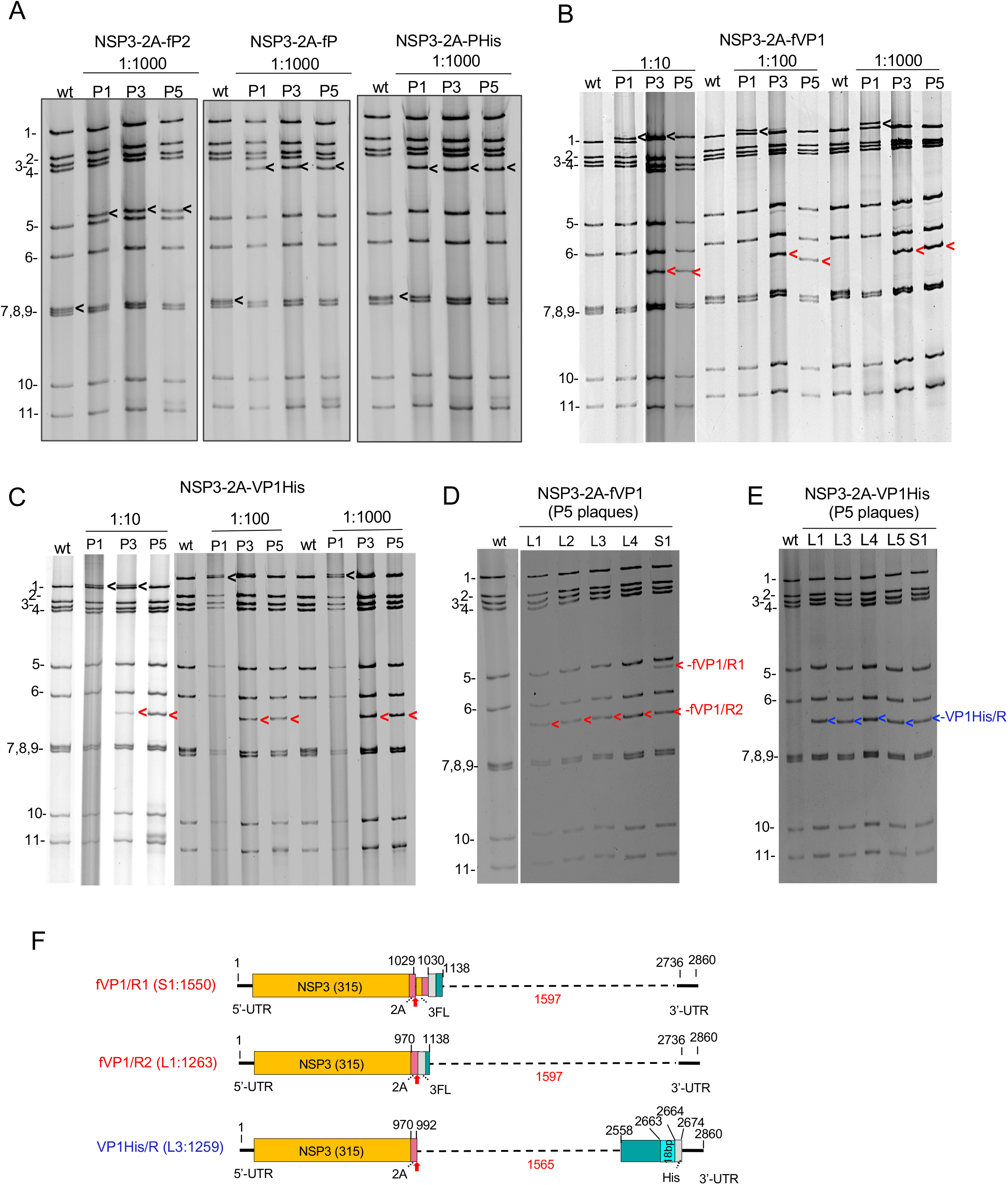
Genetic stability of rSA11 strains expressing NoV proteins. **(A-C)** rSA11 strains were serially passaged 5 times (P1 to P5) in MA104 cells, with inoculum prepared from infected cell lysates diluted 1:10, 1:100, or 1:1000 in DMEM. RNAs were recovered from infected cell lysates by Trizol extraction and analyzed by gel electrophoresis. Positions of viral genome segments are labeled. Position of modified segment 7 (NSP3) dsRNAs introduced into rSA11 strains are denoted with black arrows. **(B-C)** Novel RNAs appearing in the RNA population during serial passage are indicated with red arrows. **(D-E)** Genomic RNAs of virus isolates recovered by plaque isolation from P5 virus pools, with large (L) and small (S) plaque phenotypes. Novel (re-arranged) segment 7 RNAs are denoted with red (fVP1/R1, fVP1/R2) and blood (VP1 His/R) arrows. **(F)** Organization of re-arranged segment 7 RNAs of fVP1/R1, fVP1/R2, and VP1 His/R. Dashed lines represent sequences deleted from the starting segment 7 RNAs. Features identified include the NSP3 ORF (orange), 2A peptide (red), VP1 ORF (blue-green), 3xFLAG tag (3FL), 6xHis tag (His). Segment 7 of the VP1His/R variant contains an 18 bp sequence duplication between 2663 and 2664 (cyan). Red arrow indicates the 2A cleavage site. Nucleotide positions correspond to those of the starting non-rearranged segment 7 RNAs.

To understand the origin of the ∼1.2 kbp variant RNA, plaque isolation was used to recover five viruses each from the P5 virus pools of rSA11/NSP3-2A-fVP1 and -VP1His, four with a large (L) plaque phenotype and one with a small (S) plaque phenotype. Gel electrophoresis showed that none of the plaque isolated viruses contained the 2.8-2.9 kbp segment 7 RNAs of the unpassaged virus (Fig. 8D, E). Rather, the L1-L4 isolates of the NSP3-2A-fVP1 P5 pool contained the fVP1/R2 RNA segment (R, rearranged), whereas the S1 isolate contained both fVP1/R1 and fVP1/R2 RNA segments (Fig. 8D). Gel electrophoresis indicated that the large (L1, L3-5) and small (S1) plaque isolates of the rSA11/NSP3-2A-VP1His P5 pool contained only a single type of variant RNA: VP1His/R (Fig. 8E).

Sequence analysis revealed that the fVP1/R1 and fVP1/R2 RNAs originated from the 2.9 kbp segment 7 RNA of rSA11/NSP3-2A-fVP1 and that the VP1His/R variant RNA originated from the 2.8 kbp segment 7 RNA of rSA11/ NSP3-2A-VP1His (Fig. 8F). While the fVP1/R1 (1550 bp) and fVP1/R2 (1263 bp) RNAs contained the complete 5’-UTR and NSP3 ORF of segment 7, they lacked 1.6 kbp of the NoV VP1 coding sequence and 7 bp of the 3’-UTR. Alignment of the fVP1/R1 and fVP1/R2 sequences indicated that fVP1/R1 contained a 287-base sequence duplication comprised of residues present at the C terminus of NSP3 ORF and the 2A peptide. Comparison of the fVP1/R1 and VP1/R2 sequences suggests that the fVP1/R2 RNA may have derived from the fVP1/R1 RNA, through a secondary sequence rearrangement that resulted in the deletion of the duplicated NSP3 ORF/2A sequence (Fig. 8F). Sequence analysis showed that the VP1His/R RNA retained the complete 5’-UTR, NSP3 ORF and 3’-UTR of the segment 7 RNA. Unexpectedly, the VP1His/R RNA was found to contain two distinct alterations of its VP1 coding sequence: one, a ∼1.6 kbp deletion of the first three-fourths of the VP1 ORF and, two, an 18 bp sequence duplication of residues present near the end of the VP1 ORF (Fig. 8F). The fact that VP1 ORF of VP1His/RNA contains two alterations indicates that the segment 7 RNA of the VP1His/R variant is likely to have gone through two recombination events.

The fact that all the five virus isolates from the P5 pool of the rSA11/NSP3-2A-fVP1 and NSP3-2A-VP1His viruses contained variant 7 RNAs that were much smaller than the initial 2.8-2.9 kbp segment 7 RNAs of these viruses, suggest that smaller RNAs provide a growth advantage (Fig. 8D, E). This is consistent with a previous report suggesting that selective pressures exist during rotavirus replication that favor minimizing segment size (38). Although all three variant RNAs (fVP1/R1, fVP1/R2, and VP1His/R) contained deletions in NoV VP1 coding sequences, the RNAs all retained complete sequences for the NSP3 ORF suggesting that NSP3 may have an essential function in viral replication. Further analysis of the total population of viral RNAs in serial passaged virus may provide better insight into mechanisms governing the deletion of VP1 sequences from rSA11/NSP3-2A-VP1 viruses.

## Discussion

Through modification of the RV segment 7 RNA, we have generated recombinant SA11 viruses that express all (VP1) or portions (P, P2) of the major capsid protein of the NoV GII.4 MD145 isolate (48). Viruses of the GII.4 genotype are the most common cause of NoV illness globally (15). Given that VP1, P, and P2 contain immunodominant epitopes are targeted by neutralizing NoV antibodies (21,22), recombinant RVs expressing these proteins may be able to trigger immunoprotective responses to both RV and NoV in the immunized host. In previous studies, VLPs assembled from baculovirus-expressed VP1 were shown to induce the formation of blocking (neutralizing) antibodies in animals (49). Although we do not know whether rSA11-expressed VP1 likewise assembled into VLPs, results were obtained indicating that rSA11-expressed VP1 formed dimers and was recognized by a conformationally-dependent VP1 neutralizing antibody. Thus, rSA11-expressed VP1 may fold into native structures able to stimulate the production of neutralizing antibodies and generate immunoprotective responses in animals. Based on the dimerization and immunoprecipitation assays performed in this study, the P and P2 proteins expressed by rSA11 viruses appear not to have folded into structures that mimic the P and P2 protruding domains of the NoV capsid. However, through appropriate genetic engineering of the expressed P protein, it may be possible to generate modified forms of P that self-assemble into structures resembling P-particles and are capable of inducing NoV-specific immunoprotective responses.

Recombinant rSA11 viruses expressing VP1, P, or P2 were generated though replacement of the NSP3 ORF in the segment 7 RNA with an ORF sequentially encoding NSP3, a 3-residue flexible (GAG) linker, a 19-aa porcine teschovirus 2A translation element, and a FLAG or 6xHis tagged VP1, P, or P2 protein. Because none of the viral ORFs of the rSA11 viruses were altered except that of NSP3, these viruses are expected to produce the same complement of proteins as wild type virus. Even though NSP3 of rSA11s expressing NoV proteins contained an extra 19-aa at their C-terminus (NSP3-2A), representing remnant residues of the cleaved 2A peptide (44), the NSP3-2A product was able to dimerize. This finding is consistent with previous results showing that the NSP3 product of similarly constructed rSA11 viruses expressing fluorescent proteins also dimerizes (37). Moreover, NSP3 with 2A remnant residues also retain the ability to induce nuclear accumulation of PABP (37). The fact that NSP3 functions are not affected by remnant 2A residues may be expected given that NSP3 of group C rotavirus (RVC) is naturally expressed with a downstream 2A peptide. This allows the RVC NSP3 genome segment to express an additional protein (double-stranded RNA-binding protein, dsRBP). In our analysis of rSA11 viruses, we found that teschovirus 2A elements were efficient in promoting the expression of NoV capsid proteins as independent products. In some cases, low levels of read-through products (NSP3-2A-NoV protein) were detected, due to a failure of 2A function. In general, read-through products were more noticeable when FLAG or 6xHis antibody tags, or extra amino acids, was positioned between the 2A element and NoV protein sequence. This is consistent with earlier reports indicating that 2A activity can be affected by the nature of upstream and downstream residues (50).

The maximum amount of foreign sequence that can be inserted into the RV genome has not been determined. However, naturally-occurring RV strains have been isolated that contain 0.9-kbp sequence duplications in their segment 7 RNA, increasing the RNA’s total size to 2.0 kbp (47). In this study, generation of the rSA11/NSP3-2A-fVP1 virus required insertion of 1.8 kbp of foreign sequence into the 1.1-kbp segment 7 RNA, bring the RNA’s total size to 2.9 kbp, well in excess of segment 7 RNAs of viruses with natural sequence duplications. The total size of the genome of rSA11/NSP3-2A-fVP1 was 20.3 kbp. Although this significantly exceeds the 18.5-kbp genome of wild type SA11, larger recombinant RVs have been recovered. For example, a recombinant rSA11 virus has been generated with a 2.2-kbp segment 7 RNA that expresses the S1 protein of SARS-CoV-2 S1 (38).

Recombinant RVs with large sequence insertions have a small plaque phenotype, suggesting the extra sequence imposes a limitation on some aspect of viral growth. Interestingly, rSA11 viruses expressing NoV VP1 not only had a small plaque phenotype, but the segment 7 RNAs of these viruses were genetically unstable. During 2 to 3 rounds of serial passage, the 1.8-kbp VP1 sequence insertion in the segment 7 RNAs were lost, giving rise to variant virus populations that outcompeted the parental strains. Molecular mechanisms driving deletion of the VP1 sequences are not known, but clearly indicate that rSA11 viruses with 1.8-kbp VP1 sequence insertions are under considerable pressure either to delete the foreign sequence or to reduce the overall size of the viral genome. The instability results observed here with rSA11 viruses expressing NoV VP1 are much like those observed previously with rSA11 viruses expressing the SARS-CoV-2 S1 protein (38). In that case, the modified segment 7 RNAs of the viruses were found to delete most, if not all, of their 2.2-kbp S1 sequence insertions within a few rounds of serial passage.

In contrast to rSA11 viruses with 1.8-kbp VP1 sequence insertions, rSA11 viruses containing 1.1-kbp 3xFLAG-P sequences or 0.6-kbp 3xFLAG-P2 sequences were genetically stable over 5 rounds of serial passage (37,38). Likewise, rSA11 viruses with modified segment 7 RNAs containing sequences of the SARS-CoV-2 spike gene up to 1.5 kpb were found to be genetically stable. Thus, recombinant RVs with foreign sequence insertions in segment 7 that do not exceed 1.5 kpb may be sufficiently genetically stable for developing vaccine candidates. The coding capacity provided by 1.5 kbp of foreign sequence is sufficient to generate recombinant RVs that express the NoV P protein (∼35 kDa) or modified forms of NoV P with localization or affinity tags that may increase immunogenicity potential. For example, inclusion of affinity ligands for Fc-immunoglobulin G1 (Fc-IgG1) or for the neonatal cell surface (FcRn) receptor may help to direct expressed P protein that will promote antigen recognition and presentation (51-53).

This represents the second study illustrating the potential usefulness of RV as an expression platform for the capsid protein of another pathogenic virus. In an earlier study, through similar modification of the segment 7 RNA, we showed that it was possible to generate recombinant RVs that express regions of the SARS-CoV-2 spike protein, including the immunodominant S1 region. In unpublished work, we have also generated rSA11 viruses that express portions of the capsid proteins of astrovirus and hepatitis E virus (54, data not shown). Importantly, we have found in this study that the segment 7 RNAs of other RV strains, such as the RIX4414 (G1P[8]) vaccine strain, can be engineered to serve as expression platforms of non-rotaviral proteins. Given that reverse genetics systems have been developed for human G1P[8] and G4P[8] RVs, it seems likely that RV strains that formulates human RV strains can be developed into combination vaccines that target other enteric or mucosal viruses. With the development of reverse genetics systems against animal RV strains, it may be possible to develop similar combination vaccines for use in livestock and farm animals.

## Materials and Methods

### Cell culture

Embryonic monkey kidney (MA104) cells were grown in Dulbecco’s Modified Eagle Medium (DMEM) containing 5% fetal bovine serum (FBS) and 1% penicillin-streptomycin (55). Baby hamster kidney cells constitutively expressing T7 RNA polymerase (BHK-T7) were provided by Dr. Ulla Buchholz, Laboratory of Infectious Diseases, NIAID, NIH, and were propagated in Glasgow minimum essential media (GMEM) containing 5% heat-inactivated FBS, 10% tryptone-peptide broth, 1% penicillin-streptomycin, 2% non-essential amino acids, and 1% glutamine (56). BHK-T7 cells were grown in a medium supplemented with 2% Geneticin (Invitrogen) with every other passage.

### Plasmid construction

rSA11 viruses were prepared using the plasmids pT7/VP1SA11, pT7/VP2SA11, pT7/VP3SA11, pT7/VP4SA11, pT7/VP6SA11, pT7/VP7SA11, pT7/NSP1SA11, pT7/NSP2SA11, pT7/NSP3SA11, pT7/NSP4SA11, and pT7/NSP5SA11 [https://www.addgene.org/Takeshi_Kobayashi/] and pCMV/NP868R (36). The plasmid pT7/NSP3-2A-fUnaG was produced by fusing a DNA fragment containing the ORF for 2A-3xFL-UnaG to the 3’-end of the NSP3 ORF of pT7/NSP3SA11 using a Takara In-Fusion cloning kit. A plasmid (pUC57/MDA145_VP1) containing a full-length cDNA of the VP1 gene of the NoV GII.4 MD145-12 strain (GenBank: AY032605.1) was purchased from Genewiz. The plasmids pT7/NSP3-2A-fP2, pT7/NSP3-2A-fP, pT7/NSP3-2A-fVP1 were made by replacing the UnaG ORF in pT7/NSP3-2A-fUnaG with ORFs for the P2, P, and VP1 regions, respectively, of the NoV VP1 capsid protein, by In-Fusion cloning. The backbone of the plasmids was generated through PCR amplification of pT7/NSP3-2A-fUnaG with the primer pairs Vector_For and Vector_Rev (Table 2). DNA fragments containing P2, P, and VP1 coding sequences were amplified from pUC57/MDA145_VP1 using the primer pairs fP2_For and fP2_Rev, fP_For and fP_Rev, fVP1_For and fVP1_Rev respectively (Table 2). The plasmids pT7/NSP3-2A-VP1f, pT7/NSP3-2A-P-His, pT7/NSP3-2A-VP1-His, pT7/NSP3-2A-VP1-Th His were made similarly. The backbone of the plasmids was generated by amplifying pT7/NSP3-2A-fUnaG with the primer pairs Vector P2A_For and Vector P2A_Rev (Table 2). DNA fragments containing VP1 with a C terminal FLAG or P or VP1 with a C terminal His tag were produced through PCR amplification of pUC57/MDA145_VP1 with the primer pairs VP1-fFor and VP1-fRev, P-His_For and P-His_Rev, VP1-ThHis_For and VP1-ThHis_Rev, VP1-His_For and VP1-His_Rev respectively (Table 2). A puc19 plasmid containing a RIX/NSP3-2A-P-His insert under the control of a T7 transcription promoter (puc19/T7/ RIX/NSP3-2A-P-His) was purchased from Bio Basic Canada Inc. Transfection quality plasmids were prepared commercially (www.plasmid.com) or using Qiagen plasmid purification kits. Primers were provided by and sequences determined by EuroFins Scientific.

**Table 2.**
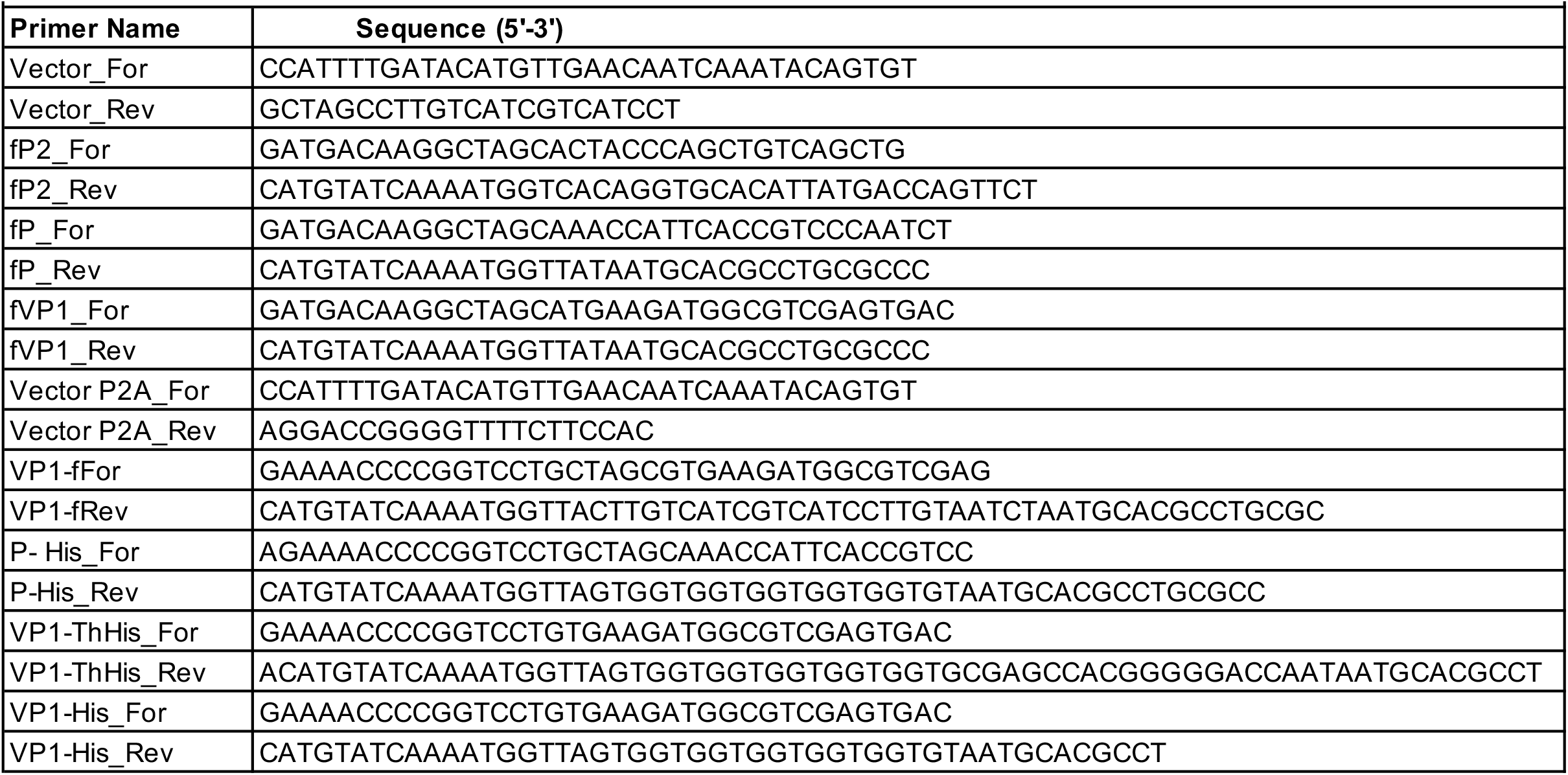
Primers used to produce pT7/NSP3-2A-NoV plasmids.

### Recombinant viruses

The reverse genetics protocol used to generate recombinant rotaviruses was described in detail previously (36,37). Briefly, BHK-T7 cell monolayers in 12-well plates were transfected with SA11 pT7 plasmids and pCMV-NP868R using Mirus TransIT-LT1 transfection reagent. Transfection mixtures contain 0.8 μg of each of the 11 pT7 plasmids except for pT7/NSP2SA11 and pT7/NSP5SA11, which were used at levels 3-fold higher. Two days after transfection, the BHK-T7 cells were over-seeded with MA104 cells and the trypsin in the medium was adjusted to a final concentration of 0.5 μg/ml. Three days later, the BHK-T7/MA104 cell mixture was freeze-thawed 3-times and the lysates were clarified by low-speed centrifugation (800 x *g*, 5 min). Recombinant viruses in clarified lysates were amplified by a single round of passage on MA104 monolayers and recovered by plaque purification (55,56). Viral dsRNAs were recovered from infected-cell lysates by TRIzol extraction, resolved by electrophoresis on 10% polyacrylamide gels in Tris-glycine buffer, detected by staining with ethidium bromide, and visualized using a BioRad ChemiDoc MP Imaging System.

### Plaque assay

Rotavirus plaque assays were performed as described before (55). To visualize plaques, cell monolayers with agarose overlays were incubated overnight with phosphate-buffered saline (PBS) containing 3.7% formaldehyde. Afterward, agarose overlays were removed, and the monolayers were stained for 3 h with a solution of 1% crystal violet dissolved in 5% ethanol. Monolayers then were rinsed with water and air-dried. Plaque images were captured using a Bio-Rad ChemiDoc imaging system and diameters were measured using ImageJ software and the results were analyzed with GraphPad Prism, version 8. Statistical significance of plaque size differences was determined using an unpaired Student’s *t*-test and included 95% confidence intervals.

### Immunoblot analysis

MA104 cells were mock-infected or infected with 5 plaque-forming units (PFU) per cell of recombinant virus and harvested at 9 h p.i. Cells were washed with cold PBS, pelleted by centrifugation (5000 x *g*, 10 min), and lysed by incubation for 30 min on ice in non-denaturing lysis buffer (300 mM NaCl, 100 mM Tris-HCl, pH 7.4, 2% Triton X-100, and 1x EDTA-free protease inhibitor cocktail [Roche Complete]). For immunoblot assays, lysates were resolved by electrophoresis on 10% polyacrylamide gels and transferred to nitrocellulose membranes. After blocking with PBS containing 5% nonfat dry milk, blots were probed with mouse monoclonal FLAG M2 (F1804, Sigma, 1:2000), mouse monoclonal His tag antibody (MCA1396GA, Bio-Rad, 1:1000), mouse 2A antibody (NBP2-59627, Novus, 1:1000), guinea pig polyclonal NSP3 (Lot 55068, 1:2000), VP6 (Lot 53963, 1:2000) antisera or rabbit monoclonal β-actin (8457S, Cell Signaling Technology (CST), 1:1000) antibody. Primary antibodies were detected using 1:10,000 dilutions of horseradish peroxidase (HRP)-conjugated secondary antibodies (goat anti-mouse IgG (CST), goat anti-guinea pig IgG (KPL), or goat anti-rabbit IgG (CST) or Alexa fluor conjugated antibody (goat anti-mouse Alexa 647 antibody (CST) in 2.5% (Carnation) nonfat dry milk. HRP signals were developed using Clarity Western ECL Substrate (Bio-Rad) and detected using a Bio-Rad ChemiDoc imaging system, whereas Alexa fluor signals were visualized directly using a Bio-Rad ChemiDoc imaging system. To evaluate the dimerization capacity of NoV proteins expressed by rSA11 viruses, cell lysates were adjusted to a final concentration of 1.5% sodium dodecyl sulfate and 3% β-mercaptoethanol and incubated for 10 min at 25°C or 95°C. Afterward, proteins in the samples were resolved by electrophoresis on 10% polyacrylamide gels and detected by immunoblot assay.

### Immunoprecipitation assay

Whole-cell lysates were prepared at 9 h p.i from MA104 cells either mock-infected or infected with rSA11 virus, as described above. Rabbit anti-NoV GII.4 monoclonal antibody [(Absolute Antibody, NVB43.9), final dilution of 1:150] was added to cell lysates. After incubation at 4°C with gentle rocking for 18 h, antigen-antibody complexes were recovered using Pierce magnetic IgA/IgG beads (ThermoScientific), resolved by gel electrophoresis, and blotted onto nitrocellulose membranes. Blots were probed with antibodies specific for FLAG (1:2000) and 6xHis tags (1:1000) to detect FLAG and 6xHis tagged VP1 proteins, respectively.

### Genetic stability of rSA11 viruses

Viruses were serially passaged five times on MA104-cell monolayers using 1:1000, 1:100, or 1:10 dilutions of infected cell lysates prepared in serum-free DMEM medium. When cytopathic effects reached completion (4-5 days), the cells were freeze-thawed three times in their own medium, and the lysates were clarified by low-speed centrifugation. dsRNAs were recovered from the clarified lysates by TRIzol extraction. The purified dsRNAs were resolved by electrophoresis on 10% polyacrylamide gels, and the bands of dsRNA were detected by ethidium bromide staining.

### Isolation and sequencing of unstable variants

Individual rSA11 variants were recovered from pools of the serially passaged virus by plaque isolation (55). The variants were amplified by a single round of passage on MA104 cells and their genomic dsRNA recovered by Trizol extraction. The full-length genome segment 7 RNAs in the samples were amplified with gene-specific primer pairs NSP3_5’UTR 5’GGCATTTAATGCTTTTCAGTG 3’ and NSP3_3’UTR 5’ GGCCACATAACGCCCCTATAG 3’, and a shorter fragment from the C terminus of NSP3 ORF to 3’UTR region was amplified with the primer pairs NSP3 C termF 5’ CATTGCACGCTTTTGATGACTTAG 3’ and NSP3_3’UTR 5’GGCCACATAACGCCCCTATAG 3’ similarly using Superscript III One-Step RT-PCR System with Platinum *Taq* DNA polymerase (Invitrogen). Amplified PCR products were resolved by electrophoresis on 0.8% agarose gels in Tris-acetate-EDTA buffer, products were gel-purified using Nucleospin gel and PCR Clean-up (Takara), and the sequences were determined by EuroFins Scientific.

### GenBank accession numbers

Segment 7 sequences in rSA11 viruses have been deposited in GenBank: wt (LC178572), NSP3-2A-NoVfP2 (MN190002), NSP3-2A-NoVfP (MN190003), NSP3-2A-NoVfVP1 (MN190004), NSP3-2A-NoV VP1f (MN201548), NSP3-2A-NoV P-His (MN201549), NSP3-2A-NoV VP1-ThHis (MN201547), NSP3-2A-NoV VP1-His (MZ562305), RIX/NSP3-2A-NoV P-His (MZ643978). See also Table 1.

## ACKNOWLEDGMENTS

Our appreciation goes out to all the members of the rotavirus research group for their support and encouragement on the project. Our thanks also go out to Ulla Buckholtz and Peter Collins (NIAID, NIH) for providing BHK-T7 cells and to Takeshi Kobayashi for making the SA11 pT7 RG plasmids available. This work was supported by funds provided by the National Institutes of Health (R21AI144881), Indiana Clinical and Translational Sciences Institute, and the Lawrence M. Blatt Endowment.

